# PreTSA: computationally efficient modeling of temporal and spatial gene expression patterns

**DOI:** 10.1101/2024.03.20.585926

**Authors:** Haotian Zhuang, Zhicheng Ji

## Abstract

Modeling temporal and spatial gene expression patterns in large-scale single-cell and spatial transcriptomics data is a computationally intensive task. We present PreTSA, a method that offers computational efficiency in modeling these patterns and is applicable to single-cell and spatial transcriptomics data comprising millions of cells. PreTSA consistently matches the results of state-of-the-art methods while significantly reducing computational time. PreTSA provides a unique solution for studying gene expression patterns in extremely large datasets.

## Main

Single-cell RNA-sequencing (scRNA-seq) and spatial transcriptomics (ST) technologies offer unprecedented opportunities to study the temporal and spatial dynamics of gene expression with single-cell or near single-cell resolution. While scRNA-seq does not directly retain or measure the order of cells in continuous biological processes such as cell development and differentiation, such ordering information can be computationally inferred using pseudotime analysis^1–6^. This allows for the examination of how a gene’s expression level changes temporally along a continuous process^7–13^. ST, on the other hand, measures both the gene expression and spatial locations of single cells or small clusters of cells, enabling the direct interrogation of spatial gene expression patterns from the observed data^14–19^.

A smooth function, such as that obtained by the generalized additive model (GAM), is often used to characterize the temporal and spatial patterns of gene expression. Smoothing helps to recover the real patterns of gene expression from noisy data and mitigates the visualization biases associated with plotting numerous data points^20^. Furthermore, a statistical model can be built upon the smooth function to identify temporally variable genes (TVGs) and spatially variable genes (SVGs) with statistical significance. GAM has been extensively used in pseudotime methods such as `Monocle`^1,3^, `TSCAN`^2^, and `Slingshot`^4^. `PseudotimeDE`^21^ further applies subsampling and permutation approaches to account for the uncertainty in pseudotime inference, thereby generating well-calibrated *p*-values for the identification of TVGs. While most existing methods for analyzing ST data, such as `SpatialDE`^22^, `SPARK`^23^, `SPARK-X`^24^, and `nnSVG`^25^, focus on identifying SVGs, GAM can also be extended to fit and visualize spatial gene expression patterns principally.

The number of cells included in scRNA-seq and ST datasets has been increasing exponentially in recent years, leading to datasets that often contain millions of cells^5,26–30^. Most existing methods, however, were developed and tested on much smaller datasets, typically comprising only thousands of cells. Consequently, they may not be computationally feasible for handling such large datasets. For example, GAM involves complex computations such as penalized regression splines and must be fitted for each gene individually, which could be computationally intensive for large datasets with thousands of genes. The repeated subsampling and permutation employed by `PseudotimeDE` further exacerbates this issue. Additionally, `nnSVG`, a method for analyzing ST data, can take more than a minute to analyze a single gene and up to a week to analyze thousands of genes, despite its ability to scale linearly.

To address this issue, we developed `PreTSA` (<u>P</u>attern <u>r</u>ecognition in <u>T</u>emporal and <u>S</u>patial <u>A</u>nalyses), a computationally efficient method designed to handle extremely large datasets. For each gene, `PreTSA` fits a regression model using B-splines, obtaining a smoothed curve that represents the relationship between gene expression values and pseudotime. Given the thousands of genes, this approach results in thousands of regression models. These models share the same design matrix, as the pseudotime values, which are specific to cells, remain consistent across different genes. Thus, `PreTSA` performs all computations related to the design matrix once, unlike existing methods such as GAM, which repeat these computations for each gene (Figure 1a). This simplification substantially reduces the computational burden, particularly for large datasets. In addition, `PreTSA` leverages efficient matrix operations in R to further enhance computational efficiency. By default, `PreTSA` employs the simplest B-spline basis without internal knots to achieve optimal computational speed. Users have the option to select a flexible mode, `PreTSA-K`, which automatically determines the number of knots in the B-spline basis (Methods). `PreTSA` utilizes the `PseudotimeDE` framework for identifying TVGs through a subsampling and permutation procedure. In addition to serving as a standalone pipeline, `PreTSA` can also be integrated into other frameworks to replace GAM and increase computational efficiency.

**Figure 1.**
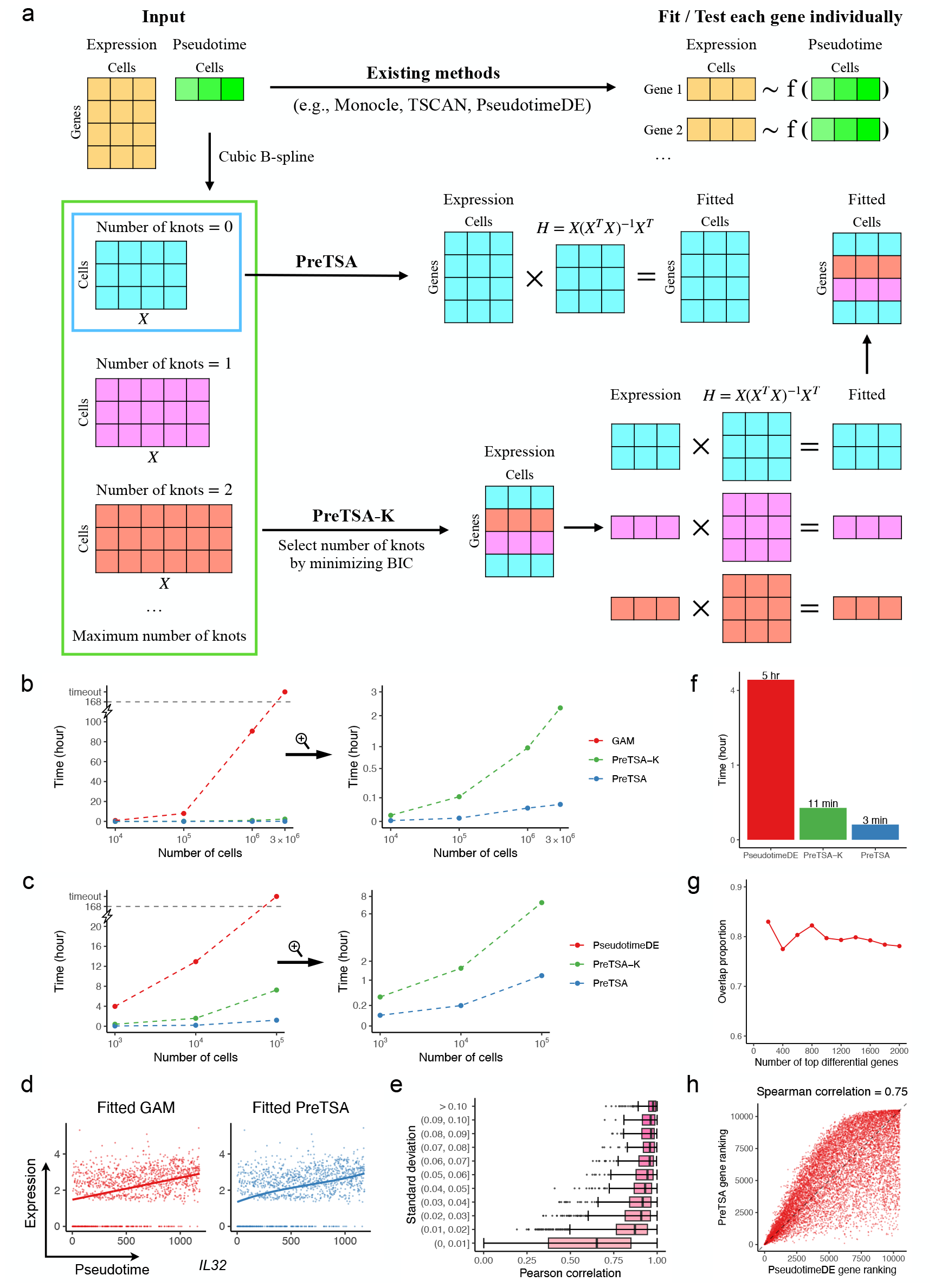
Temporal modeling of gene expression patterns by `PreTSA`. **a**, A schematic view of `PreTSA` for modeling temporal gene expression patterns. **b-c**, Computational time (y-axes) of different methods for fitting the temporal pattern (**b**) and testing TVGs (**c**) in simulated data with different numbers of cells (x-axes, log10 scale). Computational time more than one week (168 hours) is marked as timeout. The right panels are zoom in views of the left panels, with the y-axes being in square root scale. **d**, Scatterplots showing the expression of *IL32* gene (y-axis) and the pseudotime (x-axis). The curve indicates the fitted curve by GAM (left) or by `PreTSA` (right). **e**, Pearson correlations between fitted values by GAM and by `PreTSA` (x-axis), grouped by the standard deviation of fitted values by GAM (y-axis). **f**, Computational time of different methods for testing TVGs in the real scRNA-seq dataset. **g**, Overlap proportion for different numbers of top differential genes using `PreTSA` and `PseudotimeDE`. **h**, Gene rankings by `PreTSA` (y-axis) and by `PseudotimeDE` (x-axis).

We first compared the computational efficiency of `PreTSA` and GAM in modeling the temporal pattern of gene expression. We applied both methods to a simulated dataset derived from a real scRNA-seq dataset of peripheral blood mononuclear cells (PBMCs, Methods). The simulated dataset comprises 10,509 genes and ranges from thousands to millions of cells. All computations were carried out on a 3.0 GHz CPU using a single thread. `PreTSA` substantially outperformed GAM in computational efficiency (Figure 1b). For analyzing one million cells, GAM took 90.6 hours, while `PreTSA` required only 2 minutes. For three million cells, GAM failed to complete the computation within a week (168 hours), whereas `PreTSA` took merely 3 minutes. `PreTSA-K`, which involves additional steps to select the number of knots, displayed computational efficiency comparable to `PreTSA` and was still much faster than GAM. A similar disparity in computational efficiency was observed between `PreTSA` and `PseudotimeDE` in identifying TVGs (Figure 1c). For analyzing one hundred thousand cells, `PseudotimeDE` did not finish the computation within a week, while `PreTSA` took 1.2 hours. These results suggest that `PreTSA` is the only method capable of completing computations within a reasonable timeframe for large datasets.

We next compared the temporal gene expression patterns modeled by `PreTSA` and GAM in the original PBMC dataset. Figure 1d displays the expression and fitted curves for the example gene, *IL32*, known to be associated with the T cell activation process^31^. The curves fitted by `PreTSA` and GAM are highly similar, both indicating linearly increasing trends. Figure 1e presents the Pearson correlation coefficients (PCCs) between the curves fitted by `PreTSA` and GAM for each gene, where the median PCC exceeds 0.9 for genes with standard deviations of GAM-fitted curves larger than 0.01. `PreTSA-K` demonstrates highly similar findings (Supplementary Figure 1a-b). Further, we compared the results of `PreTSA` and `PseudotimeDE` in identifying TVGs. Notably, in this smaller dataset comprising thousands of cells, `PseudotimeDE` required five hours to complete, whereas `PreTSA` took only three minutes (Figure 1f). Genes were ordered increasingly by the *p*-values from `PreTSA` and `PseudotimeDE` for identifying TVGs, respectively. The overlap in the top-ranked genes identified by both methods is approximately 80% (Figure 1g). The Spearman correlation coefficient for the overall gene rankings by both methods is as high as 0.75 (Figure 1h). `PreTSA-K` yielded highly similar results (Supplementary Figure 1c-d). These findings indicate that `PreTSA` produces results highly consistent with those of existing methods while substantially increasing computational efficiency.

`PreTSA` can also be applied to ST data by extending the one-dimensional B-spline bases to two dimensions using tensor products (Figure 2a). Since the spatial locations of cells are directly measured, `PreTSA` uses an F-test to identify SVGs, without accounting for additional variance as in the case of identifying TVGs. Similarly, GAM can be extended with tensor products. Using a real 10x Visium dataset of the human heart, we constructed a simulated dataset with 11,953 genes and thousands to millions of spatial spots. Note that although the number of spots in 10x Visium datasets is relatively small, the latest ST technologies, such as 10x Visium HD, 10x Xenium, MERSCOPE, and NanoString CosMx, can easily produce a much larger number of spots or cells. Thus, the simulated dataset reflects the different scenarios one may encounter in practice. `PreTSA` again demonstrates superior computational efficiency over GAM (Figure 2b). For analyzing one million spots, GAM took 68.3 hours, while `PreTSA` required only 7 minutes. For analyzing two million spots, GAM failed to complete the computation within a week, while `PreTSA` took just 8 minutes. For identifying SVGs, `PreTSA` and `SPARK-X` are the only methods that can handle a million spots within an hour (Figure 2c). Methods such as `SpatialDE`, `SPARK`, `SPARK-G`, and `nnSVG` failed to complete the computation within a week even for a hundred thousand spots, and GAM was unable to handle a million spots within a week.

**Figure 2.**
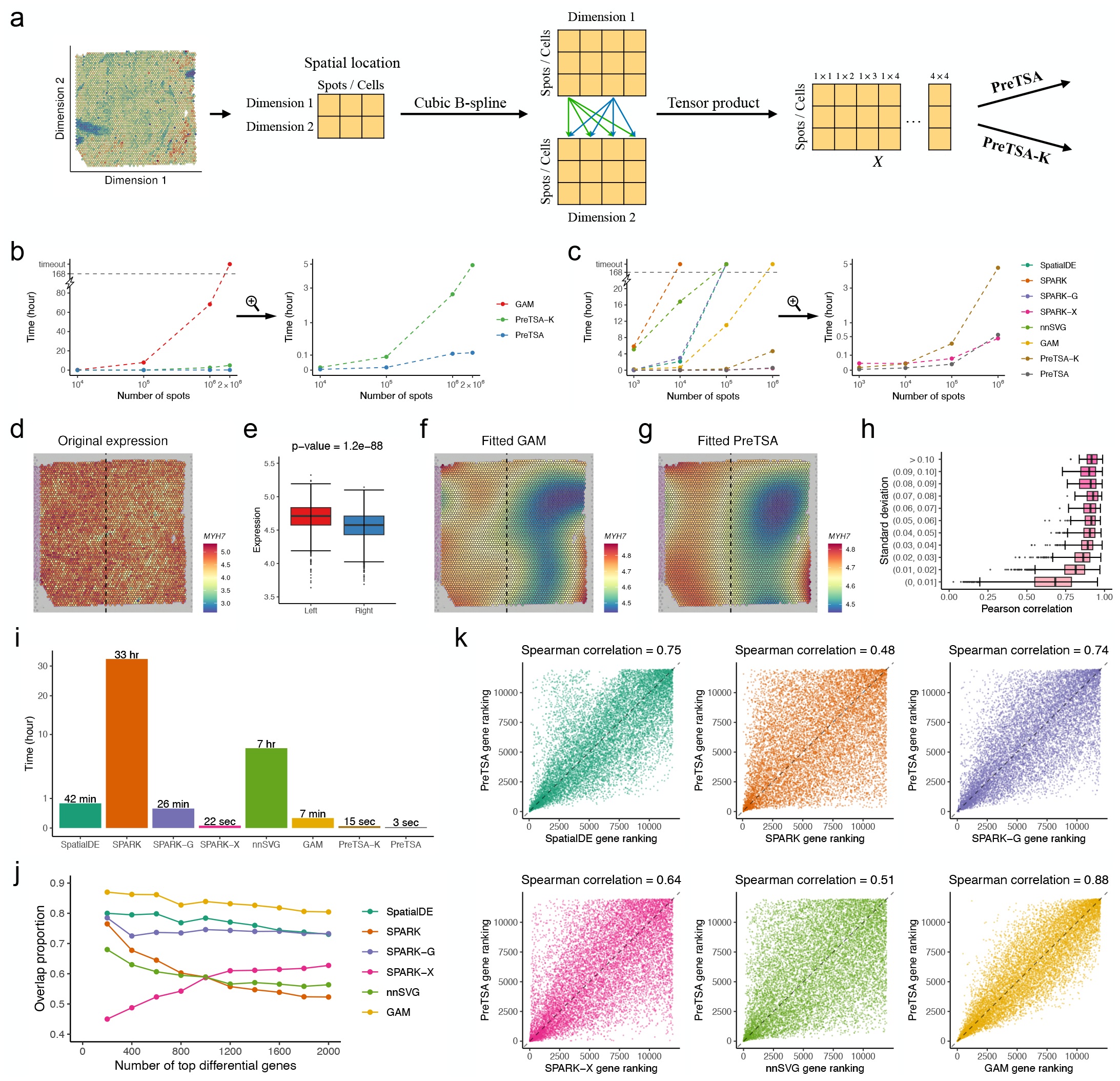
Spatial modeling of gene expression patterns by `PreTSA`. **a**, A schematic view of `PreTSA` for modeling spatial gene expression patterns. **b-c**, Computational time (y-axes) of different methods for fitting the spatial pattern (**b**) and testing SVGs (**c**) in simulated data with different numbers of spots (x-axes, log10 scale). Computational time more than one week (168 hours) is marked as timeout. The right panels are zoom in views of the left panels, with the y-axes being in square root scale. **d**, Original expression of *MYH7* gene in Visium dataset. **e**, Boxplot showing the distribution of gene expression in the left and right spatial regions, separated by the dotted vertical line in **d**. *p*-value is obtained by Wilcoxon rank-sum test. **f-g**, Spatial expression pattern of *MYH7* fitted by GAM (**f**) and by `PreTSA` (**g**). **h**, Pearson correlations between fitted values by GAM and by `PreTSA` (x-axis), grouped by the standard deviation of fitted values by GAM (y-axis). **i**, Computational time of different methods for testing SVGs in the real spatial dataset. **j**, Overlap proportion for different numbers of top differential genes using `PreTSA` and other methods. **k**, Gene rankings by `PreTSA` (y-axis) and by other methods (x-axis).

While `PreTSA` and `SPARK-X` have comparable computational efficiency, `SPARK-X` lacks the ability to fit the spatial pattern of gene expression, which is crucial for understanding complex ST data. Figure 2d shows the expression of the example gene *MYH7* in the original Visium heart dataset. *MYH7* plays a critical role in the functionality of cardiac muscle fibers in the human heart^32^. Although it is challenging to visually discern the spatial pattern of the gene, the expression level of *MYH7* is significantly higher in the left part of the tissue compared to the right (Figure 2e). This distinction can only be elucidated by the spatial gene expression patterns fitted by GAM and `PreTSA`, which are highly similar (Figure 2f-g). The median PCC between surfaces fitted by `PreTSA` and GAM exceeds 0.85 for genes with standard deviations of GAM-fitted surfaces larger than 0.01 (Figure 2h). `PreTSA-K` demonstrates highly similar findings (Supplementary Figure 2a-b). These results indicate that `PreTSA` is the only method capable of efficiently fitting spatial gene expression patterns in large datasets.

Finally, we compared the results of identifying SVGs using `PreTSA` and other existing methods in the original Visium heart dataset. Both `PreTSA` and `SPARK-X` completed computations within a minute, whereas methods such as `SPARK` and `nnSVG` required several hours (Figure 2i). The genes were ordered increasingly by the *p*-values from different methods for identifying SVGs. The top-ranked genes identified by `PreTSA` show the most similarity to those identified by GAM, `SpatialDE`, and `SPARK-G`, with an overlap proportion of around 80% (Figure 2j). The Spearman correlation coefficients for the overall gene rankings also reached at least 0.74 (Figure 2k). The results from `PreTSA` are less consistent with those from `SPARK`, `SPARK-X`, and `nnSVG`. However, `SPARK`, `SPARK-X`, and `nnSVG` are more distinct from any other existing methods, whereas GAM, `SpatialDE`, and `SPARK-G` are more similar to each other (Supplementary Figure 2c). `PreTSA-K` demonstrated highly similar results (Supplementary Figure 2d-e). While it is hard to assess the quality of identified SVGs due to the lack of a gold standard and varying definitions of SVGs^33,34^, these results suggest that `PreTSA` is consistent with a group of existing methods for identifying SVGs.

In summary, we have demonstrated that `PreTSA` substantially improves computational efficiency in modeling temporal and spatial gene expression patterns, while producing results that are consistent with state-of-the-art methods, such as GAM, `PseudotimeDE`, and `SpatialDE`. `PreTSA` offers a unique solution for understanding the temporal and spatial dynamics of gene expression in large single-cell and ST datasets.

## Methods

### PreTSA model for temporal gene expression patterns

#### Inputs

`PreTSA` requires as input a numeric matrix of library-size-normalized and log-transformed gene expression values, along with a numeric vector of pseudotime values for single cells. The gene expression matrix can be derived from any standard processing pipeline for scRNA-seq data^35–37^. Pseudotime values can be obtained from any pseudotime inference method^1–5^. Users also have the option to replace the pseudotime values with any other one-dimensional numeric values of interest.

#### Fitting temporal gene expression patterns

Denote **Y** as the *m* × *n* gene expression matrix, with *m* genes and *n* cells, and *y*_*i j*_ as the expression level of gene *i* in cell *j*. Denote *t* _*j*_ as the pseudotime value for cell *j*. For each gene *i*, we model its expression values along pseudotime as a functional curve:

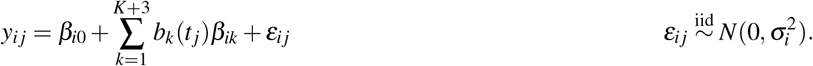

Here, *b*_1_(*t*), …, *b*_*K*+3_(*t*) represent the *K* + 3 cubic B-spline basis functions, where *K* is the number of equidistant internal knots used to define the cubic B-spline bases. The method for selecting *K* will be discussed later. The parameters *β*_*i*0_, …, *β*_*i*(*K*+3)_ and 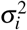 are all unknown and will be estimated using the least squares method.

For gene *i*, the regression model can be written in matrix form as follows: **Y**_*i*_ = **X*β*** _*i*_ + ***ε***_*i*_, where

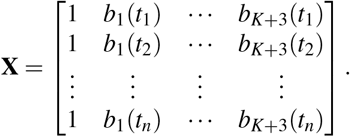

Note that the hat matrix **H** = **X**(**X**^*T*^ **X**)^−1^**X**^*T*^ is shared across all genes. Consequently, we can apply the hat matrix directly to the original gene expression matrix to obtain the fitted gene expression matrix **Ŷ** as follows:

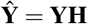

Here *ŷ*_*i j*_ represents the fitted gene expression value of gene *i* in cell *j*.

`PreTSA` sets *K* = 0 for all genes. `PreTSA-K`, an extended model of `PreTSA`, enables the automatic selection of *K* for each gene by minimizing the Bayesian Information Criterion (BIC). For a given *K* and a given gene *i*, 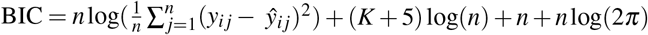.

By default, `PreTSA-K` chooses *K* from the set of integers ranging from 0 to 10.

#### Testing for temporally variable genes (TVGs)

To test whether the expression level of gene *i* is temporally variable, or equivalently, cannot be represented by a horizontal line along the pseudotime axis, we define the following null and alternative hypotheses:

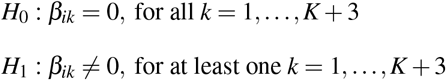

`PreTSA` uses the F-statistics:

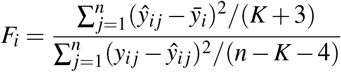

Here 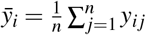.

To account for the uncertainty inherent in computationally inferring pseudotime, we apply the same strategy used in `PseudotimeDE`^21^. We first subsample 80% of the cells, a proportion suggested by `PseudotimeDE`^21^, and then apply the same pseudotime inference method that was used for the original dataset to this subsampled dataset to obtain the pseudotime values. These values are subsequently randomly permuted. Following this, the fitting approach described earlier is applied to the subsampled and permuted dataset to obtain null F-statistics. This entire process is repeated 100 times to generate 100 null F-statistics. To enhance numerical accuracy, a Gamma distribution is fitted to these 100 null F-statistics using the R package `fitdistrplus` (version 1.1.11). The *p*-value is calculated as the probability of the tail of the fitted Gamma distribution being greater than *F*_*i*_. All *p*-values are then adjusted for multiple testing using the Benjamini-Hochberg (BH) procedure^38^.

### PreTSA model for spatial gene expression patterns

#### Inputs

`PreTSA` requires two inputs: a numeric matrix of library-size-normalized and log-transformed gene expression values, and a numeric matrix of 2-dimensional spatial locations for cells or spatial spots. The gene expression matrix can be obtained through any standard processing pipeline for ST data^35,39,40^. The 2-dimensional spatial location information is usually directly available from the ST dataset.

#### Fitting spatial gene expression patterns

Denote **Y** as the *m* × *n* gene expression matrix, with *m* representing number of genes and *n* representing number of cells or spots, and *y*_*i j*_ as the expression level of gene *i* in cell or spot *j*. Denote **S** _*j*_ = (*s* _*j*1_, *s* _*j*2_) as the 2-dimensional spatial location of cell or spot *j*.

For each gene *i*, we model its expression values across spatial locations as a functional surface:

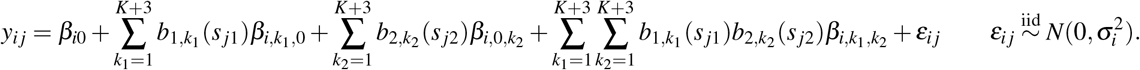

Here, *b*_*d*,1_(*s*), …, *b*_*d,K*+3_(*s*) represent the *K* + 3 cubic B-spline basis functions for each dimension (*d* = 1, 2). The parameters *β*_*i*0_, *β*_*i*,1,0_, …, *β*_*i,K*+3,0_, *β*_*i*,0,1_, …, *β*_*i*,0,*K*+3_, *β*_*i*,1,1_, …, *β*_*i,K*+3,*K*+3_, and 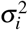 are all unknown and will be estimated using the least squares method.

For gene *i*, the regression model can be written in matrix form as follows: **Y**_*i*_ = **X*β*** _*i*_ + ***ε***_*i*_, where

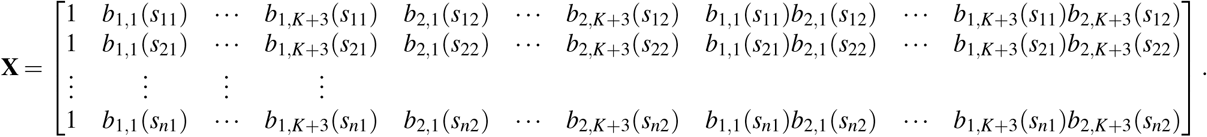

Similar to the case of modeling the temporal gene expression pattern, we can directly apply the hat matrix to the original gene expression matrix to obtain the fitted gene expression matrix **Ŷ**:

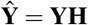

Here *ŷ*_*i j*_ represents the fitted gene expression value of gene *i* in cell or spot *j*.

`PreTSA` chooses *K* = 0 for all genes. `PreTSA-K` chooses the optimal *K* using the same BIC procedure as the case of fitting temporal gene expression pattern. For a given *K* and a given gene *i*, 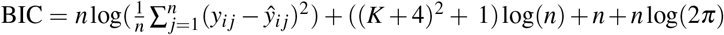.

By default, `PreTSA-K` chooses *K* from the set of integers ranging from 0 to 5.

#### Testing for spatially variable genes (SVGs)

To test whether the expression level of gene *i* varies spatially, similar to the case of testing for TVGs, we define the following null and alternative hypotheses:

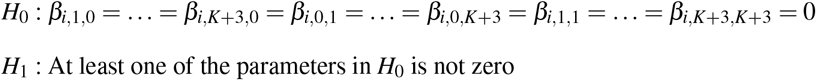

`PreTSA` uses the F-statistics:

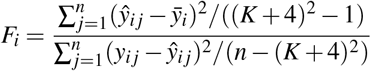

which follows an exact F-distribution with ((*K* + 4)^2^ − 1, *n* − (*K* + 4)^2^) degrees of freedom under *H*_0_.

Here 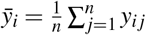. All *p*-values are then adjusted for multiple testing using the BH procedure^38^.

### Data processing

#### PBMC scRNA-seq dataset

The processed PBMC dataset was downloaded using the R package `SeuratData` (version 0.2.2). It included the log-normalized gene expression matrix, principal components (PCs) and cell type annotations. `TSCAN`^2^ (version 2.0.0) was performed using the top 10 PCs linking naive CD4 T cells and memory CD4 T cells, using the R function `guided_tscan()`. The direction of the pseudotime is from naive CD4 T cells to memory CD4 T cells. Genes that are expressed in fewer than 5 cells within the pseudotime trajectory were filtered out. A total of 10,509 genes and 1,180 cells were obtained.

#### Visium spatial human heart dataset

The Visium dataset of a human heart tissue was downloaded from the 10x website https://www.10xgenomics.com/resources/datasets/human-heart-1-standard-1-1-0. `Seurat`^35^ (version 4.3.0) was used to process the data. Mitochondrial genes and genes expressed in fewer than 50 spots were filtered out. A total of 11,953 genes and 4,247 spots were obtained. `SCTransform`^41^ (version 0.4.1) with default settings was used to normalize and log-transform the raw count data.

### Simulation datasets

For PBMC scRNA-seq dataset, we randomly sampled 1 thousand, 10 thousand, 100 thousand, 1 million, and 3 million cells with replacement to create simulated datasets. The gene expression profiles of these cells are the same as in the original dataset. The pseudotime values of these cells are assigned as a sequence of consecutive integers, starting from one and continuing through to the total number of cells.

For Visium spatial human heart dataset, we randomly sampled 1 thousand, 10 thousand, 100 thousand, 1 million, and 2 million spots with replacement to create simulated datasets. The gene expression profiles of these spots are the same as in the original dataset. The spatial locations of these spots are assigned as all possible combinations of two sequences of consecutive integers. The first sequence ranges from 1 to *i*, and the second sequence ranges from 1 to *j*, for each pair of values (*i, j*) specified. In this simulation, the specific pairs used are (25, 40), (80, 125), (250, 400), (800, 1250), (1000, 2000), respectively.

### Competing methods

For modeling temporal gene expression patterns, GAM was implemented using the R package `mgcv` (version 1.8.41) with the formula *y*∼ *s*(*t, k* = 3), same as `TSCAN`^2^. `PseudotimeDE`^21^ (version 1.0.0) with the R function `runPseudotimeDE(model` `= “gaussian”)` was used for testing TVGs. GAM and `PseudotimeDE` used the same normalized and log-transformed gene expression matrix as used by `PreTSA`.

For modeling spatial gene expression patterns, GAM was implemented using the R package `mgcv` (version 1.8.41) with the formula *y*∼ *te*(*s*_1_, *s*_2_), where *s*_1_ and *s*_2_ are vectors representing spatial locations in two dimensions, respectively. To test SVGs, GAM performs an asymptotic chi-squared likelihood ratio test where the full model fits the gene expression across spatial locations and the null model considers the gene expression as a constant across spatial locations. In addition, we applied `SpatialDE`^22^ (version 1.1.3), `SPARK`^23^ (version 1.1.1), `SPARK-G`^23^ (version 1.1.1), `SPARK-X`^24^ (version 1.1.1), and `nnSVG`^25^ (version 1.5.8) for testing SVGs. The R package `reticulate` (version 1.32.0) was used to call the Python package `SpatialDE` within R. `SPARK` and `SPARK-X` used the gene expression count values as the input. All other methods used the same log-normalized expression values by `SCTransform` as used by `PreTSA`.

### Overlap proportion

To quantify the similarity of the top-ranked genes identified by different methods, overlap proportion is defined as the proportion of top-ranked genes identified by both methods. Specifically, denote *A* and *B* as the set of top *L* genes identified by two methods. | *A*| = |*B*|= *L*, where |.| is the cardinality of a set. The overlap proportion is defined as 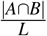. The overlap proportion was calculated for different choices of *L*.

## Availability of data and materials

All datasets used in the study are publicly available. The R package `PreTSA` with a detailed user manual is publicly available at https://github.com/haotian-zhuang/PreTSA.

## Acknowledgments

The project was supported by the National Institutes of Health under Award Number U54AG075936.

## Author contributions

Z.J. conceived the study. H.Z. developed the method and conducted the analysis. H.Z. and Z.J. wrote the manuscript.

## References

1. Trapnell, C. et al. The dynamics and regulators of cell fate decisions are revealed by pseudotemporal ordering of single cells. Nat. biotechnology 32, 381–386 (2014).

2. Ji, Z. & Ji, H. Tscan: Pseudo-time reconstruction and evaluation in single-cell rna-seq analysis. Nucleic acids research 44, e117–e117 (2016).

3. Qiu, X. et al. Reversed graph embedding resolves complex single-cell trajectories. Nat. methods 14, 979–982 (2017).

4. Street, K. et al. Slingshot: cell lineage and pseudotime inference for single-cell transcriptomics. BMC genomics 19, 1–16 (2018).

5. Cao, J. et al. The single-cell transcriptional landscape of mammalian organogenesis. Nature 566, 496–502 (2019).

6. Saelens, W., Cannoodt, R., Todorov, H. & Saeys, Y. A comparison of single-cell trajectory inference methods. Nat. biotechnology 37, 547–554 (2019).

7. Chen, Z. et al. Tcf-1-centered transcriptional network drives an effector versus exhausted cd8 t cell-fate decision. Immunity 51, 840–855 (2019).

8. Wang, W. et al. Single-cell transcriptomic atlas of the human endometrium during the menstrual cycle. Nat. Medicine 26, 1644–1653 (2020).

9. Zhang, S. et al. Longitudinal single-cell profiling reveals molecular heterogeneity and tumor-immune evolution in refractory mantle cell lymphoma. Nat. communications 12, 2877 (2021).

10. Caushi, J. X. et al. Transcriptional programs of neoantigen-specific til in anti-pd-1-treated lung cancers. Nature 596, 126–132 (2021).

11. Gearty, S. V. et al. An autoimmune stem-like cd8 t cell population drives type 1 diabetes. Nature 602, 156–161 (2022).

12. Chen, C.-W. et al. Adaptation to chronic er stress enforces pancreatic β -cell plasticity. Nat. communications 13, 4621 (2022).

13. Hou, W. et al. A statistical framework for differential pseudotime analysis with multiple single-cell rna-seq samples. Nat. communications 14, 7286 (2023).

14. Liu, Y. et al. High-spatial-resolution multi-omics sequencing via deterministic barcoding in tissue. Cell 183, 1665–1681 (2020).

15. Maynard, K. R. et al. Transcriptome-scale spatial gene expression in the human dorsolateral prefrontal cortex. Nat. neuroscience 24, 425–436 (2021).

16. Kuppe, C. et al. Spatial multi-omic map of human myocardial infarction. Nature 608, 766–777 (2022).

17. Sampath Kumar, A. et al. Spatiotemporal transcriptomic maps of whole mouse embryos at the onset of organogenesis. Nat. Genet. 55, 1176–1185 (2023).

18. Peirats-Llobet, M. et al. Spatially resolved transcriptomic analysis of the germinating barley grain. Nucleic Acids Res. 51, 7798–7819 (2023).

19. Weber, L. M. et al. The gene expression landscape of the human locus coeruleus revealed by single-nucleus and spatially-resolved transcriptomics. eLife 12, RP84628 (2024).

20. Hou, W. & Ji, Z. Unbiased visualization of single-cell genomic data with scubi. Cell reports methods 2 (2022).

21. Song, D. & Li, J. J. Pseudotimede: inference of differential gene expression along cell pseudotime with well-calibrated p-values from single-cell rna sequencing data. Genome biology 22, 124 (2021).

22. Svensson, V., Teichmann, S. A. & Stegle, O. Spatialde: identification of spatially variable genes. Nat. methods 15, 343–346 (2018).

23. Sun, S., Zhu, J. & Zhou, X. Statistical analysis of spatial expression patterns for spatially resolved transcriptomic studies. Nat. methods 17, 193–200 (2020).

24. Zhu, J., Sun, S. & Zhou, X. Spark-x: non-parametric modeling enables scalable and robust detection of spatial expression patterns for large spatial transcriptomic studies. Genome biology 22, 1–25 (2021).

25. Weber, L. M., Saha, A., Datta, A., Hansen, K. D. & Hicks, S. C. nnsvg for the scalable identification of spatially variable genes using nearest-neighbor gaussian processes. Nat. communications 14, 4059 (2023).

26. Cao, J. et al. A human cell atlas of fetal gene expression. Science 370, eaba7721 (2020).

27. Tian, Y. et al. Single-cell immunology of sars-cov-2 infection. Nat. biotechnology 40, 30–41 (2022).

28. Braun, E. et al. Comprehensive cell atlas of the first-trimester developing human brain. Science 382, eadf1226 (2023).

29. Zhang, M. et al. Molecularly defined and spatially resolved cell atlas of the whole mouse brain. Nature 624, 343–354 (2023).

30. Yao, Z. et al. A high-resolution transcriptomic and spatial atlas of cell types in the whole mouse brain. Nature 624, 317–332 (2023).

31. Goda, C. et al. Involvement of il-32 in activation-induced cell death in t cells. Int. immunology 18, 233–240 (2006).

32. Witjas-Paalberends, E. R. et al. Mutations in myh7 reduce the force generating capacity of sarcomeres in human familial hypertrophic cardiomyopathy. Cardiovasc. research 99, 432–441 (2013).

33. Heumos, L. et al. Best practices for single-cell analysis across modalities. Nat. Rev. Genet. 1–23 (2023).

34. Charitakis, N. et al. Disparities in spatially variable gene calling highlight the need for benchmarking spatial transcriptomics methods. Genome Biol. 24, 209 (2023).

35. Hao, Y. et al. Integrated analysis of multimodal single-cell data. Cell 184, 3573–3587 (2021).

36. Wolf, F. A., Angerer, P. & Theis, F. J. Scanpy: large-scale single-cell gene expression data analysis. Genome biology 19, 1–5 (2018).

37. Amezquita, R. A. et al. Orchestrating single-cell analysis with bioconductor. Nat. methods 17, 137–145 (2020).

38. Benjamini, Y. & Hochberg, Y. Controlling the false discovery rate: a practical and powerful approach to multiple testing. J. Royal statistical society: series B (Methodological) 57, 289–300 (1995).

39. Dries, R. et al. Giotto: a toolbox for integrative analysis and visualization of spatial expression data. Genome biology 22, 1–31 (2021).

40. Righelli, D. et al. Spatialexperiment: infrastructure for spatially-resolved transcriptomics data in r using bioconductor. Bioinformatics 38, 3128–3131 (2022).

41. Hafemeister, C. & Satija, R. Normalization and variance stabilization of single-cell rna-seq data using regularized negative binomial regression. Genome biology 20, 296 (2019).

