## Supplementary Figure for "PreTSA: computationally efficient modeling of temporal and spatial gene expression patterns"

### Supplementary materials

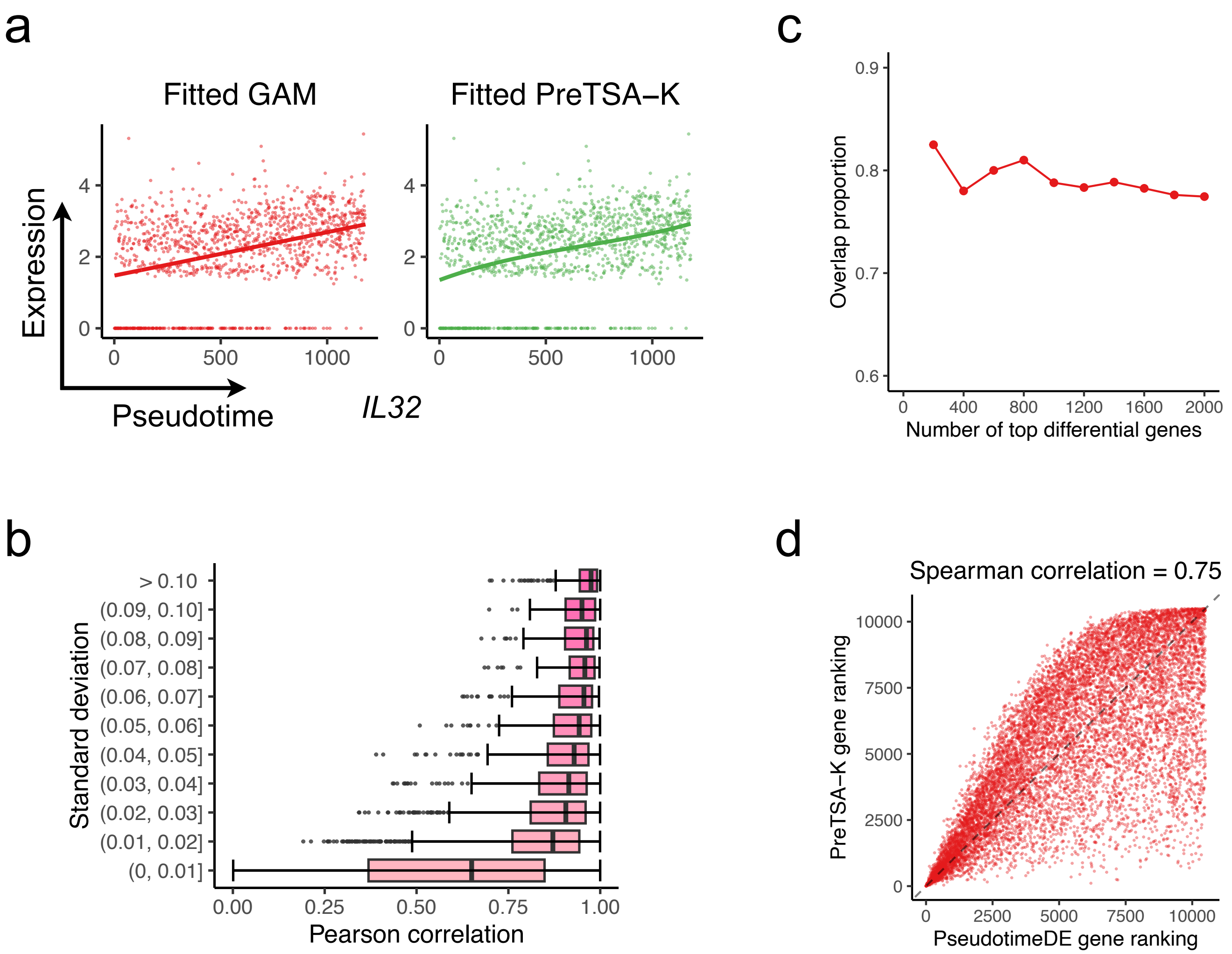


**Figure S1.** **a**, Scatterplots showing the expression of *IL32* gene (y-axis) and the pseudotime (x-axis). The curve indicates the fitted curve by GAM (left) or by `PreTSA−K` (right). **b**, Pearson correlations between fitted values by GAM and by `PreTSA−K` (x-axis), grouped by the standard deviation of fitted values by GAM (y-axis). **c**, Overlap proportion for different numbers of top differential genes using `PreTSA−K` and `PseudotimeDE`. **d**, Gene rankings by `PreTSA−K` (y-axis) and by `PseudotimeDE` (x-axis).

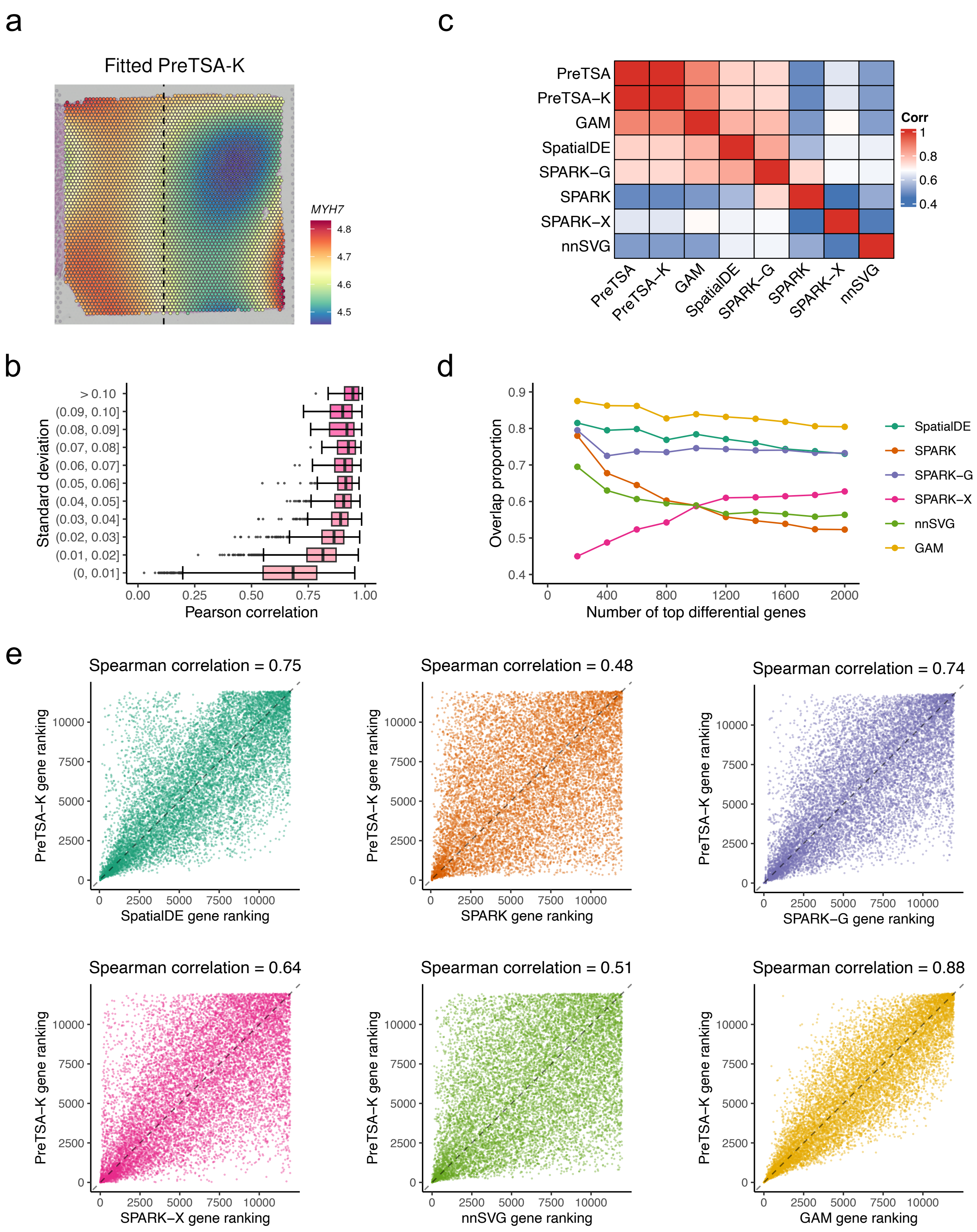


**Figure S2.** **a**, Spatial expression pattern of *MYH7* fitted by `PreTSA-K`. **b**, Pearson correlations between fitted values by GAM and by `PreTSA-K` (x-axis), grouped by the standard deviation of fitted values by GAM (y-axis). **c**, Spearman correlations for the overall gene rankings between different methods. **d**, Overlap proportion for different numbers of top differential genes using `PreTSA-K` and other methods. **e**, Gene rankings by `PreTSA-K` (y-axis) and by other methods (x-axis).
